# Trikafta rescues F508del-CFTR by tightening specific phosphorylation-dependent interdomain interactions

**DOI:** 10.1101/2024.11.20.624197

**Authors:** Guangyu Wang

## Abstract

Trikafta effectively corrects the thermal and gating defects associated with the F508del mutation, the most common cause of cystic fibrosis, even at physiological temperatures. However, the exact correction pathway is still unclear. Here, noncovalent interactions among two transmembrane domains (TMD1 and TMD2), the regulatory (R) domain and two nucleotide binding domains (NBD1 and NBD2) were analyzed. The thermal stability of NBD1 was also evaluated through its tertiary constrained noncovalent interaction networks or thermoring structures. The results demonstrated that Trikafta binding to flexible TMD1 and TMD2 rearranged their interactions with the R domain upon phosphorylation, coupling tightened cytoplasmic TMD1-TMD2 interactions to tightened Mg/ATP-dependent NBD1-NBD2 dimerization, which stabilized NBD1 above human body temperature. In essence, while the F508 deletion primarily causes a thermal defect in NBD1, leading to a gating defect at the TMD1-TMD2 interface, Trikafta allosterically reverses these effects. These mechanistic insights into the precise correction pathway of this misfolded channel facilitate optimizing cystic fibrosis treatment. (155 words)

**Key points:**

- Trikafta binding to flexible TMD1 and TMD2 tightened their cytoplasmic interactions.
- Tight cytoplasmic TMD1-TMD2 interactions primed the specific binding of the dynamic phosphorylated S813 site to the TMD1/TMD2/NBD1 interfaces.
- The tight binding of the S813 site to the TMD1/TMD2/NBD1 interfaces strengthened NBD1-NBD2 dimerization which stabilizes NBD1.

## 1. INTRODUCTION

Cystic fibrosis (CF) is the most common fatal genetic disease in the Caucasian population and is also prevalent in other human populations. On the other hand, secretory diarrhea is the leading cause of infant mortality worldwide. A key issue in both of these devastating diseases is a protein called Cystic Fibrosis Transmembrane conductance Regulator (CFTR) with activity that is either too low or too high. CFTR belongs to the family of ATP-binding cassette (ABC) transporters but functions as a phosphorylation-dependent and ATP-gated anion (chloride or bicarbonate) channel. It regulates the salt-water balance of the lung, intestine, pancreas, and sweat once expressed on the cell plasma membrane in epithelial tissues around the human body (1–4). To date, more than 2000 mutations in CFTR have been found to cause CF (http://www.genet.sickkids.on.ca/cftr<u>; CFTR2</u> <u>database:</u> https://cftr2.org). The most common mutation (70 % of CF cases) is ΔF508, which has both gating and thermal defects, leading to airway bacterial infection and respiratory failure (4–13).

A cryo-electron microscopy (cryo-EM) structure of dephosphorylated human CFTR (PDB, 5UAK) reveals that it starts with a lasso motif (1–37) and is composed of five well-known domains in the following order: transmembrane domain 1 (TMD1) (38–383), nucleotide binding domain 1 (NBD1) (384–645), the regulatory (R) domain (646–843), transmembrane domain 2 (TMD2) (844–1172), and nucleotide binding domain 2 (NBD2) (1173–1436). Both TMD1 and TMD2 line an anion conduction pathway, each consisting of six transmembrane (TM) helices that extend to the intracellular loops (ICLs). The arrangement of TM1, 2, 3 and 6 forming a bundle with TM10 and 11, while the remaining TM helices build another bundle, allows for domain swapping. This enables NBD1 to primarily connect with TMD2 via ICL4 and NBD2 to mainly relate to TMD1 via ICL2 (14). When the R domain is phosphorylated by protein kinase A (PKA) for Mg/ATP-mediated NBD1-NBD2 dimerization, the hydrolysis-deficient E1371Q mutant is open with distinct NBD1-ICL4 and NBD2-ICL2 interactions (15, 16).

In contrast, the most common cystic fibrosis-causing ΔF508 mutant primarily destabilizes NBD1 and nearby interdomain interactions (17–21). Even if a type I corrector like lumacaftor (VX-809) is bound to TMD1 in the phosphorylated and ATP-bound conformation to secure the channel expression on the cell surface, NBD1 folding remains flexible, resembling a “molten globule” until the folding corrector tezacaftor (VX-661), the folding and gating modulator elexacaftor (VX-445) and the channel activity potentiator ivacaftor (VX-770) are combined in Trikafta (branded as Kaftrio in Europe) and bound to TMD1 and TMD2. This combination therapy completely restores ΔF508 to an NBD-dimerized conformation with ATP fully occluded at both the consensus and degenerate sites (22). Although Trikafta synergistically rescued the gating and thermal defects of ΔF508-CFTR at 37 °C (23–27), much less is known about the exact correction pathway of the most flexible NBD1.

On the other hand, the thermostability of isolated hNBD1 can be measured using differential scanning calorimetry (DSC) or differential scanning fluorimetry (DSF). It can be characterized by a melting temperature (T_m_), at which half of the protein is unfolded. It can also be defined by a melting temperature threshold (T_m,th_) for initial heat unfolding when set at 2.0 % of maximal spectral changes (9,11, 28–29). However, directly measuring NBD1’s thermostability with either biophysical approach becomes impossible in the presence of various interdomain interactions when it is not isolated from the full-length hCFTR. Therefore, an alternative tool is necessary to assess the effects of adjacent interdomain interactions on NBD1’s thermostability and to unveil the rescue pathway of the thermal defect of ΔF508 CFTR by Trikafta even within such a complex multi-domain protein.

Recently, a novel and highly sensitive graphical method has been developed to analyze the structural thermo-stability and to predict activity temperatures of proteins including enzymes (30–37). Based on their high-resolution 3D structures, the tertiary noncovalent interaction networks can be constrained by a grid thermodynamic model as the thermoring structures to stabilize each interaction along the single polypeptide. Each thermoring is denoted as Grid_s_, where “s” represents the grid size to regulate the intensity of the least-stable noncovalent interaction within this grid. It can be defined by the shortest alternative path or the minimal free amino acid side chains to link two ends of this least-stable interaction along the single polypeptide chain and other noncovalent interactions. When the biggest thermoring is identified, the melting temperature threshold (T_m,th_) to unfold the least-stable noncovalent interaction within the biggest thermoring along the single peptide can be calculated.

Since the matched theoretical and experimental T_m,th_ values of isolated human NBD1 (hNBD1) upon different structural perturbations can account for why the thermostability of NBD1 is sensitive to ATP and the disordered regulatory insert (RI) (405–436) (37), this graphical analysis can be extended to predict the T_m,th_ values of NBD1 and to evaluate the effects of the interdomain interactions on NBD1’s thermostability within the full-length hCFTR. In addition, the systematic thermal instability (T_i_) or flexibility can be evaluated by the ratio of the total grid sizes to the total noncovalent interactions along the same polypeptide chain.

The results demonstrate that Trikafta could rescue both gating and thermal defects of ΔF508 by coupling tightened cytoplasmic TMD1-TMD2 interactions to tightened Mg/ATP-mediated NBD1-NBD2 dimerization via the specific binding of the dynamic phosphorylated S813 site of the R domain to the ICL1-ICL4 interface, which is a target of curcumin (38). This coupling pathway, combined with other therapeutic strategies such as curcumin, not only improves CFTR thermal stability but also shows promise for long-term therapeutic benefits in managing cystic fibrosis. Thus, thermodynamic modeling can be utilized to understand drug action mechanisms and update protein correction strategies.

## 2. RESULTS

### 2.1 NBD1-ICL4 interactions fail to increase the melting threshold of hNBD1

The isolated hNBD1-Δ(RE, RI) (PDB, 2PZE) is a single peptide spanning from S386 to G646 with a segment deleted from 405 to 436 (28). In contrast, the dephosphorylated hNBD1 in full-length CFTR (PDB, 5UAK) has disordered RI (403–438) (14). Therefore, in the absence of Mg/ATP-mediated NBD1-NBD2 dimerization, it is interesting to compare their melting temperature thresholds and evaluate the effect of NBD1-ICL4 interactions alone on the thermal stability of hNBD1.

In the dephosphorylated state, NBD1 is separated from NBD2 by the R domain (14). Without Mg/ATP bound, NBD1 had 18 intra-domain non-covalent interactions via amino acid side chains such as seven H-bonds, four salt bridges and seven π interactions (Figures 1a & Table S1). Most of these noncovalent interactions formed smaller thermorings. Notably, a smaller Grid_5_ had a 5-residue size to control the F508-R560 and Y517-Y563 π bridges via a thermoring from F508 to Y512, Y517, Y563, R560 and back to F508. In addition, the side chain of D567 H-bonded with the backbone –NH of R487 while the backbone –NH of L568 H-bonded with the backbone –CO of R487, forming the smallest Grid_0_. In contrast, some H-bonds generated bigger thermorings. For example, when the side chain of S495 H-bonded with the side chain of R553, a bigger Grid_18_ had an 18-residue size to control the S495-R553 H-bond via a thermoring from S495 to F508, R560, R553, and back to S495. Thus, despite a total of 18 noncovalent interactions along the peptide chain from S384 to M645, a total of 71 grid sizes still resulted in an unusually high systematic thermal instability (T_i_) of 3.94 (Table 1). That may be the reason why the crystal X-ray structure of isolated hNBD1-Δ(RE, RI) is unavailable in the absence of Mg/ATP (28).

**Figure 1.**
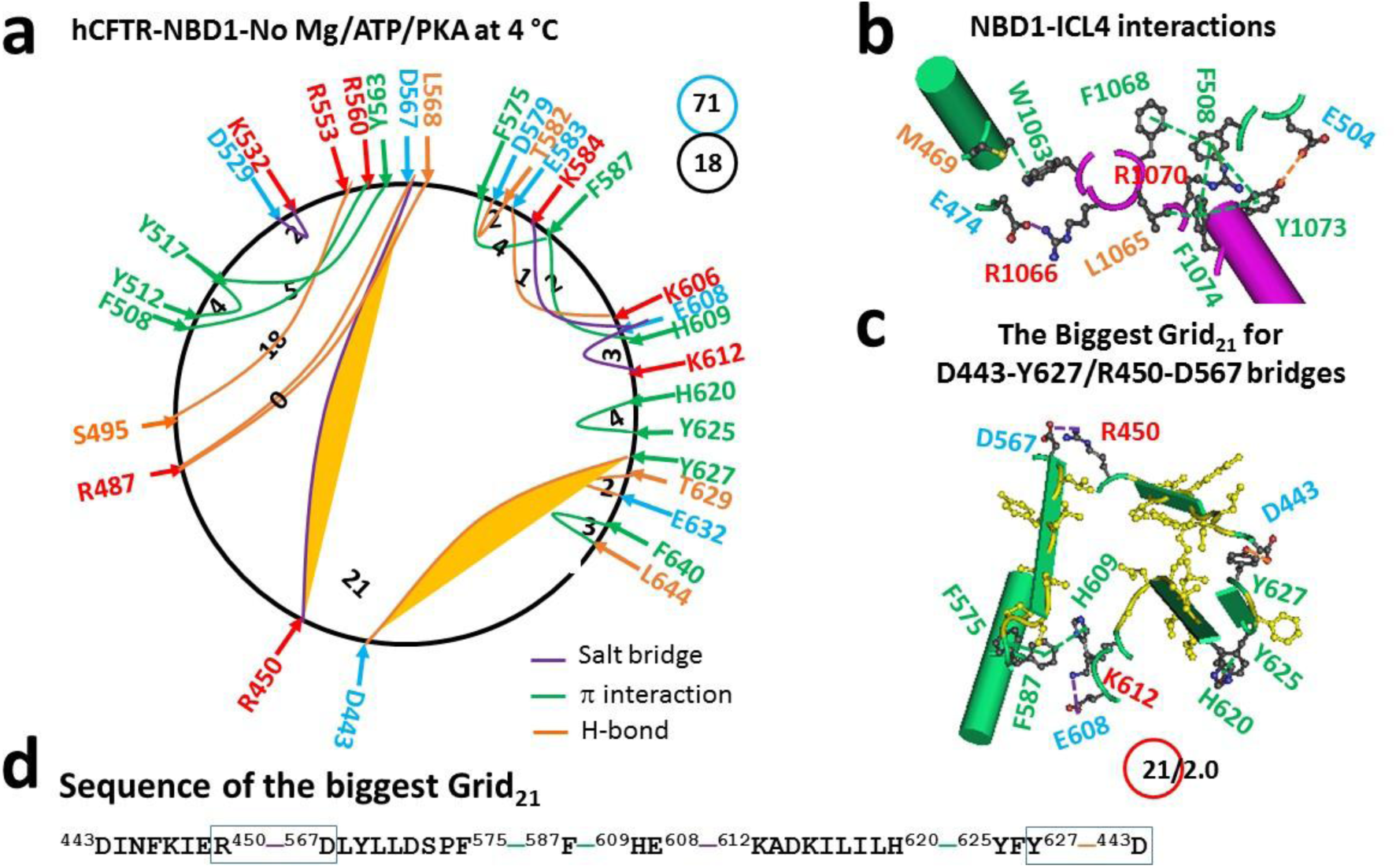
The thermoring structures of dephosphorylated hCFTR with F508 in the closed state at 4 °C. (**a**) The grid-like noncovalently interacting mesh network based on the cryo-EM structure of hCFTR with F508 in the absence of Mg/ATP/PKA at 4 °C (PDB ID, 5UAK, 3.87 Å). Salt bridges, H-bonds and π interactions are colored purple, orange, and green, respectively. The black numbers label the constrained grid sizes needed to control the least-stable noncovalent interactions in the grids. The least-stable D443-Y627 and R450-D567 bridges in the biggest Grid_21_ are highlighted. The total grid sizes and the total grid size-controlled noncovalent interactions along the single peptide chain of NBD1 from E384 to M645 are shown in cyan and black circles, respectively. (**b**) Noncovalent interactions at the NBD1/ICL4 interface. (**c**) The structure of the biggest Grid_21_ with a 21-residue size to control the least-stable D443-Y627 and R450-D567 bridges. The grid size and the equivalent basic H-bonds for the least-stable noncovalent interaction are shown in and near a red circle. (**d**) The sequence of the biggest Grid_21_ to control the least-stable D443-Y627 and R450-D567 bridges in the blue box.

**Table 1.** Grid thermodynamic model-based new parameters of NBD1 in hCFTR constructs.

| Construct | hCFTR |  |  |  |
| --- | --- | --- | --- | --- |
| PDB ID | 5UAK | 6MSM | 8EIQ | 8EIG |
| F508 | + | + | - | - |
| E1371Q | - | + | + | + |
| Mg/ATP (10 mM)/PKA | - | + | + | + |
| Sampling temperature, °C | 4 | 4 | 4 | 4 |
| Tight NBD dimerization | - | + | + | - |
| Normal Mg<sup>2+</sup> site |  | + | - | - |
| Name of the biggest grid | Grid<sub>21</sub> | Grid<sub>10</sub> | Grid<sub>11</sub> | Grid<sub>14</sub> |
| Grid size (s) | 21 | 10 | 11 | 14 |
| # of energetically equivalent basic H-bonds (n) controlled by Grid<sub>s</sub> | 2.0 | 1.6 | 1.5 | 1.3 |
| Total non-covalent interactions (N) | 18 | 40 | 36 | 31 |
| Total grid sizes (S), a.a. | 71 | 82 | 65 | 73 |
| Systematic thermal instability (T<sub>i</sub>) | 3.94 | 2.05 | 1.81 | 2.35 |
| Calculated T<sub>m</sub>, °C | **32** | **50** | **47** | **39** |
| Measured threshold T<sub>m</sub>, °C* | **32** | **48** | **40** | **40** |
| References for measured T<sub>m</sub> | (9) | (9, 28) | (9) | (9) |

Note: \* measured thresholds are based on the isolated hNBD1-Δ(RE, RI). The experimental melting threshold (T<sub>m,th</sub>) was established at 2.0 % of ΔF<sub>max</sub> or ΔC<sub>Pmax</sub>. The comparative thresholds are highlighted in bold.

In addition to the intra-domain noncovalent interactions, there were also noncovalent interactions between NBD1 and ICL4 (Figure 1b). These interactions included a salt bridge between E474 and R1066, an H-bond between E504 and Y1073, and seven π interactions such as M469-W1063, L1065-F1074-Y1073, F1068-F508-Y1073/F1074 and R1070-F508. Notably, F508, Y1073 and F1074 created the smallest triangle thermoring to stabilize the NBD1-ICL4 interface. In this case, it is necessary to examine whether these interfacial interactions affect the melting temperature threshold (T_m,th_) of NBD1.

When D443 in the N-terminal region H-bonded with Y627 in the C-terminal region of NBD1, the biggest Grid_21_ structure was created (Figure 1c). This structure had a 21-residue size to be responsible for stabilizing the least-stable D443-Y627 and R450-D567 bridges via a thermoring from D443 to R450, D567, F575, F587, H609, E608, K612, H620, Y625, Y627, and back to D443 (Figure 1c-d). These two H-bonds, when energetically equivalent to 2.0 basic H-bonds, had a calculated melting temperature threshold (T_m,th_) of 32 °C, matching the melting threshold of isolated hNBD1-Δ(RE, RI) as measured by DSC (Table 1) (9). Therefore, the presence of stronger NBD1-ICL4 interactions may not increase the melting temperature threshold of NBD1 in the absence of bound Mg/ATP.

### 2.2 Dimerization of NBD1-NBD2 increases the melting threshold of NBD1 more than NBD1-NBD1 dimerization

When hCFTR is phosphorylated for Mg/ATP-dependent NBD1-NBD2 dimerization, the NBD1 (PDB, 6MSM) exhibits a disordered segment from 638 to 646 with the unstructured RI shifting from 403-438 to 410-434 (16). Despite this minor alteration in the primary sequence structure, the tertiary noncovalent structure of NBD1 remains comparable to that of the isolated hNBD1-Δ(RE, RI) when bound by Mg/ATP for NBD dimerization (Figures 2a & Table S2) (37).

**Figure 2.**
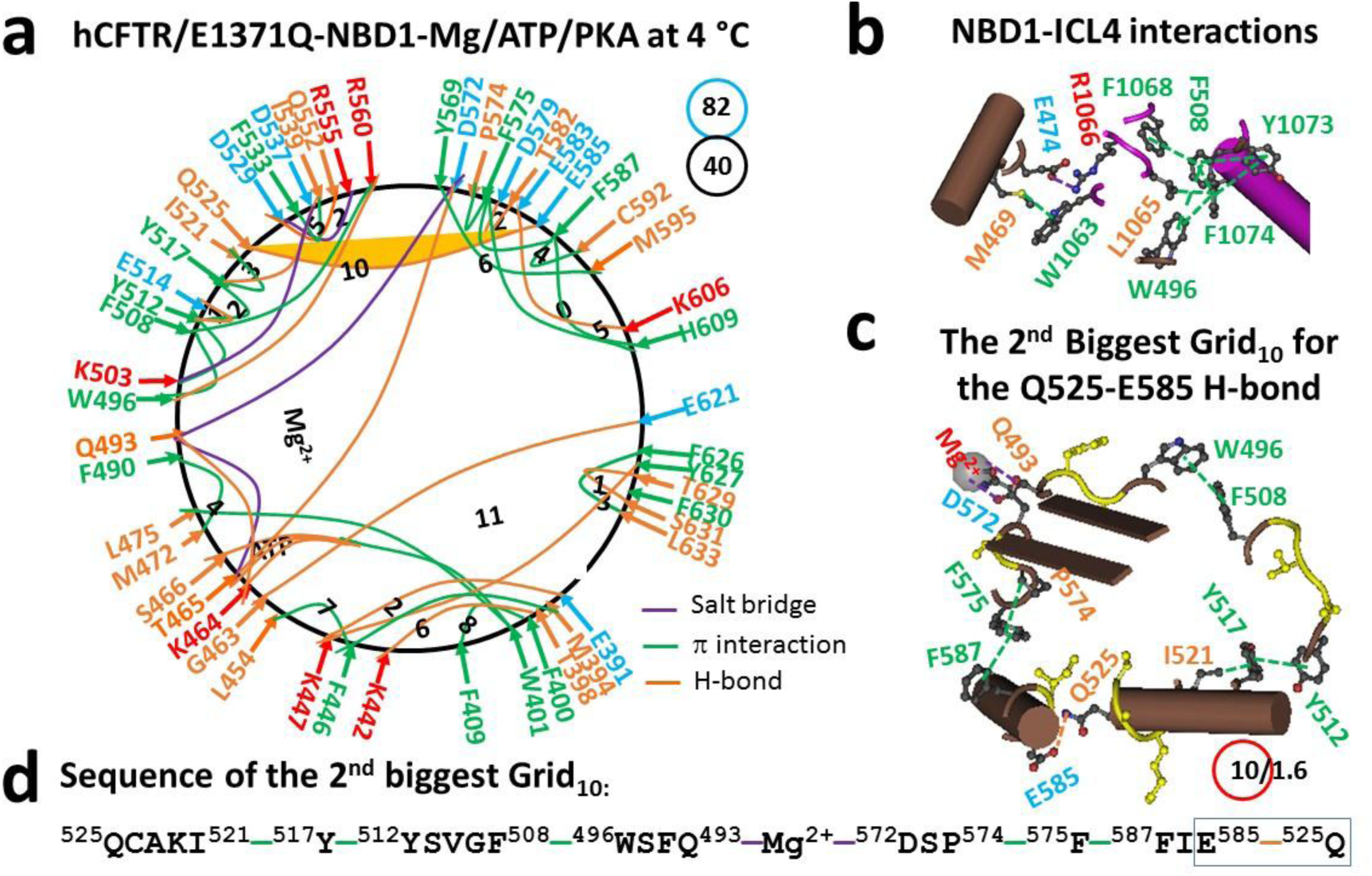
The thermoring structures of phosphorylated hCFTR/E1371Q with F508 in the open state at 4 °C. (**a**) The grid-like noncovalently interacting mesh network based on the cryo-EM structure of hCFTR/E1371Q with F508 in the presence of Mg/ATP/PKA at 4 °C (PDB ID, 6MSM, 3.2 Å). Salt bridges, H-bonds and π interactions are colored purple, orange, and green, respectively. The constrained grid sizes required to control the least-stable noncovalent interactions in the grids are labeled with black numbers. The least-stable Q525-E585 H-bond in the 2nd biggest Grid_10_ is highlighted. The total grid sizes and the total grid size-controlled noncovalent interactions along the single peptide chain of NBD1 from E384 to Q637 are shown in cyan and black circles, respectively. (**b**) Noncovalent interactions at the NBD1/ICL4 interface. (**c**) The structure of the 2nd biggest Grid_10_ with a 10-residue size to control the least-stable Q525-E585 H-bond. The grid size and the equivalent basic H-bonds for the least-stable noncovalent interaction are shown in and near a red circle. (**d**) The sequence of the 2nd biggest Grid_10_ to control the least-stable Q525-E585 H-bond in the blue box.

Firstly, the Mg/ATP binding site remained intact. For instance, T465, Q493 and D572 formed a stable triangle via Mg^2+^. Meanwhile, ATP connected W401, K464, T465, Q493, and S466. Additionally, W401 engaged in a π interaction with L475. However, the T388-T390, W401-S466, T460-ATP and R487-D567 H-bonds, as well as the F446-Y627 and H484-L568 π interactions were disrupted. Instead, the E391-K447 and T398-K442 H-bonds and the M394-F446-L454 and F400-F409 π interactions were observed.

Secondly, there were some common noncovalent bridges present near F508. These included the W496-F508 and Y512-Y517 π interactions, as well as the W496-R560, Y512-E514, Y517-E537 and D529-Q552 H-bonds and the D529-R555 salt bridge. However, the I507-Y563 π interaction, as well as the N505-R560 H-bond and the R518-E527 salt bridge were replaced with the K503-Y512, F508-R560, Y517-I521, and F533-I539 π interactions.

Thirdly, common noncovalent interactions were also observed near the Mg^2+^ site. For example, P574-F575-F587 and H609-F587/F575 π interactions. The only difference was that the D565-K598 salt bridge was substituted by the Q525-E585 H-bond, together with the disruption of the T604-H609 H-bond and the appearance of the Y569-M595 and F587-C592 π bridges.

Fourthly, in the C-terminal region of NBD1 only the F626-L633 π interaction and the T629-S631 H-bond were intact. The H620-F640-Y625 π interactions were absent, as well as the D639-S641/S642 H-bonds.

Most importantly, although most of the NBD1-ICL4 interactions were maintained upon R domain phosphorylation with the exception of the replacement of the E504-Y1073 H-bond with the L1065-F1074 π interaction (Figure 2b), the primary biggest Grid_13_ between N and C-termini changed to Grid_11_, along with a calculated melting temperature threshold (T_m,th_) of 52 °C to govern both K447-Y627 and G463-E621 H-bonds. Furthermore, the second biggest Grid_10_ was present along with the least-stable Q525-E585 H-bond (Figure 2a). It had a 10-residue size to control the new H-bond via a thermoring from Q525, I521, Y517, Y512, F508, W496, Q493, Mg^2+^, D572, P574, F575, F587, E585 and back to Q525 (Figures 2c-d). With 1.6 equivalent basic H-bonds sealing it, the calculated T_m,th_ was about 50 °C, higher than the melting threshold of 48 °C measured by DSC for the isolated hNBD1-Δ(RE, RI) homodimer (Table 1) (9, 28). Meanwhile, the systematic thermal instability (T_i_) of NBD1 dramatically decreased from 3.94 to 2.05 along with a total of 40 non-covalent bridges and 82 grid sizes (Table 1). Therefore, NBD1, when dimerized with NBD2, had a higher T_m,th_ compared to when it dimerizes with itself.

### 2.3 Dimerization of NBD1-NBD2 also increases the melting threshold of NBD1 lacking F508 more than NBD1-NBD1 dimerization does

When F508 was deleted to disrupt the W496-F508-R560 π interactions, in the presence of Trikafta, the second biggest Grid_10_ was dissociated allosterically. This also led to the disruption of the Q525-E585 H-bond (Figures 3a & Table S3). As a result, the H-bond moved from E391-K447 or T398-K442 to N396-D443 near the same Mg/ATP site along with a new salt bridge between R487 and D567, new H-bonds between W401 and S466, K442 and S623, as well as L453 and D614.

**Figure 3.**
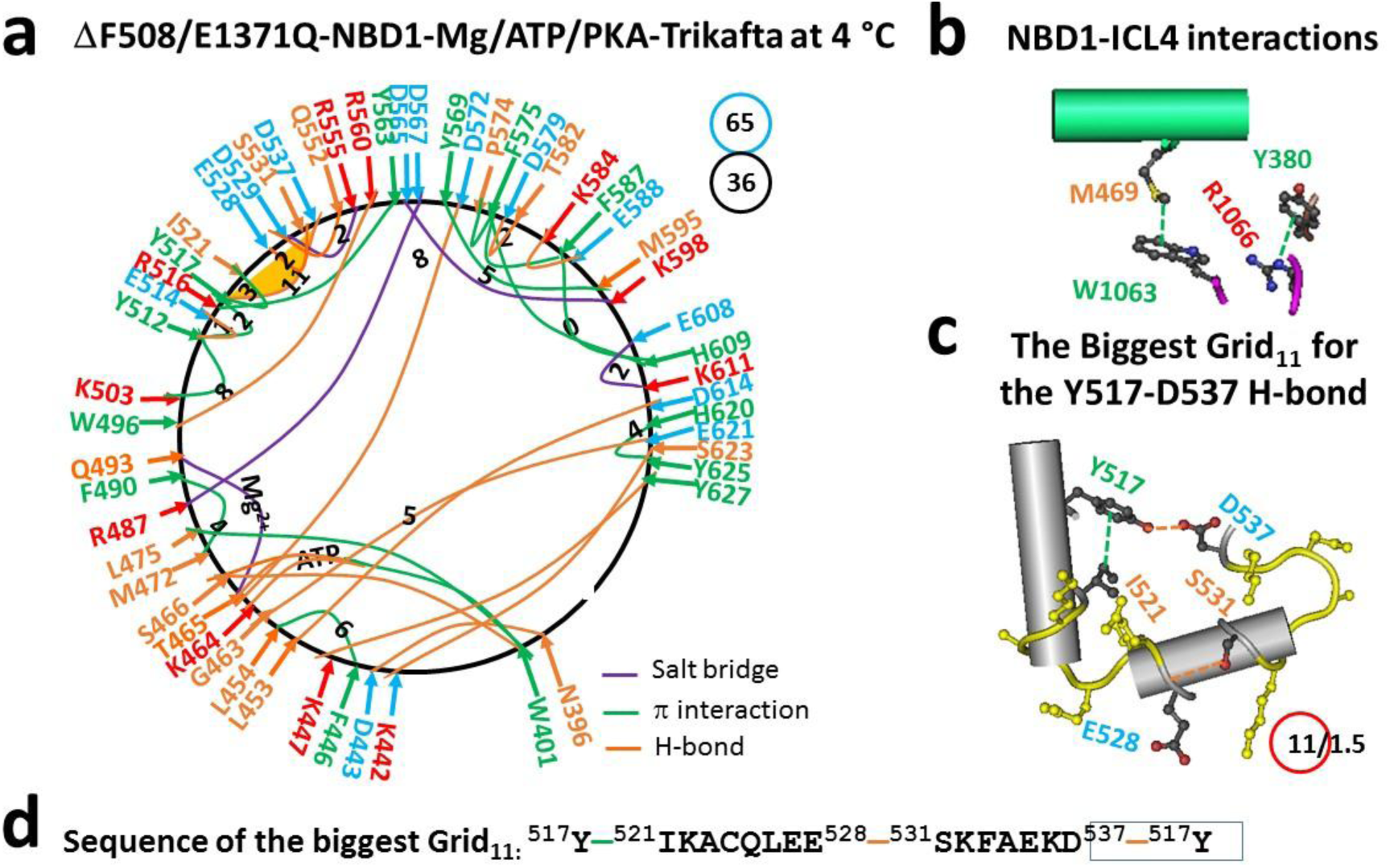
The thermoring structures of phosphorylated hCFTR/E1371Q/ΔF508 with Trikafta bound in the open state at 4 °C. (**a**) The grid-like noncovalently interacting mesh network based on the cryo-EM structure of hCFTR/E1371Q/ΔF508 with Trikafta bound in the presence of Mg/ATP/PKA at 4 °C (PDB ID, 8EIQ, 3.0 Å). Salt bridges, H-bonds and π interactions are colored purple, orange, and green, respectively. The constrained grid sizes required to control the least-stable noncovalent interactions in the grids are labeled with black numbers. The least-stable Y517-D537 H-bond in the biggest Grid_11_ is highlighted. The total grid sizes and the total grid size-controlled noncovalent interactions along the single peptide chain of NBD1 from L383 to L636 are shown in cyan and black circles, respectively. (**b**) Noncovalent interactions at the NBD1/ICL4 interface (Y380 is at the TMD1-NBD1 linker). (**c**) The structure of the biggest Grid_11_ with an 11-residue size to control the least-stable Y517-D537H-bond. The grid size and the equivalent basic H-bonds for the least-stable noncovalent interaction are shown in and near a red circle. (**d**) The sequence of the biggest Grid_11_ to control the least-stable Y517-D537H-bond in the blue box.

When this conformational change transmitted to the C-terminal region of NBD1, the F626-L633 π interaction and the T629-S631 H-bond were disconnected. Meanwhile, a new H620-Y625 π interaction were produced. Overall, when the total number of noncovalent interactions decreased from 40 to 36 along the peptide chain from E391 to Q637, the total grid sizes also decreased from 82 to 65 (Figure 3a). Thus, the systematic thermal instability (T_i_) or flexibility of NBD1 decreased from 2.05 to 1.81 along with the presence of Trikafta (Table 1).

On the other hand, the removal of F508 also disrupted most of the NBD1-ICL4 interactions except for the M469-W1063 π interaction. Additionally, the E474-R1066 salt bridge was replaced with the Y380-R1066 π interaction (Figure 3b). Despite these weakened interfacial interactions between NBD1 and ICL4, the biggest Grid_11_ was present in the α-subdomain. With an 11-residue size controlling the least-stable Y517-D537 H-bond which was equivalent to 1.5 basic H-bonds, the calculated melting threshold for the thermoring from Y517 to I521, E528, S531, D537, and back to Y517 was about 47 °C, which was higher than the melting threshold of 40 °C of the isolated (F508del)hBVD1-Δ(RE, RI) homodimer measured by DSC (Table 1) (9). Hence, the NBD1-NBD2 dimerization in the presence of Trikafta also increased the melting threshold of NBD1 without F508 by 7 °C compared to NBD1-NBD1. The next question is how Trikafta achieves this when binding to TMD1 and TMD2, rather than NBD1, the NBD1-NBD2 interface or the NBD1-ICL4 interface.

### 2.4 Tight NBD1-NBD2 dimerization primes the higher melting threshold of NBD1

A previous study demonstrated that two isolated NBD1 peptides can form a head-to-tail homodimer in the presence of Mg/ATP when the RE and RI regions are deleted (28). However, the noncovalent interactions between two NBD1 peptides are limited to the Mg/ATP site. On one side of the interface, the Walker A half-site shows extensive interactions along the phosphates, Mg^2+^ and the Walker A residues such as K464, T465 and S466. On the other side, the signature LSGGQ half-site also interacts with the phosphates through H-bonds (28).

However, when NBD1 and NBD2 dimerized, more noncovalent interactions were found between them regardless of the presence of F508del and Trikafta. In addition to the traditional interactions between the phosphates and the Walker A/B half-site or the signature LSHGH/ LSGGQ half-site (39), some new noncovalent interactions were also observed nearby (Figure 4).

**Figure 4.**
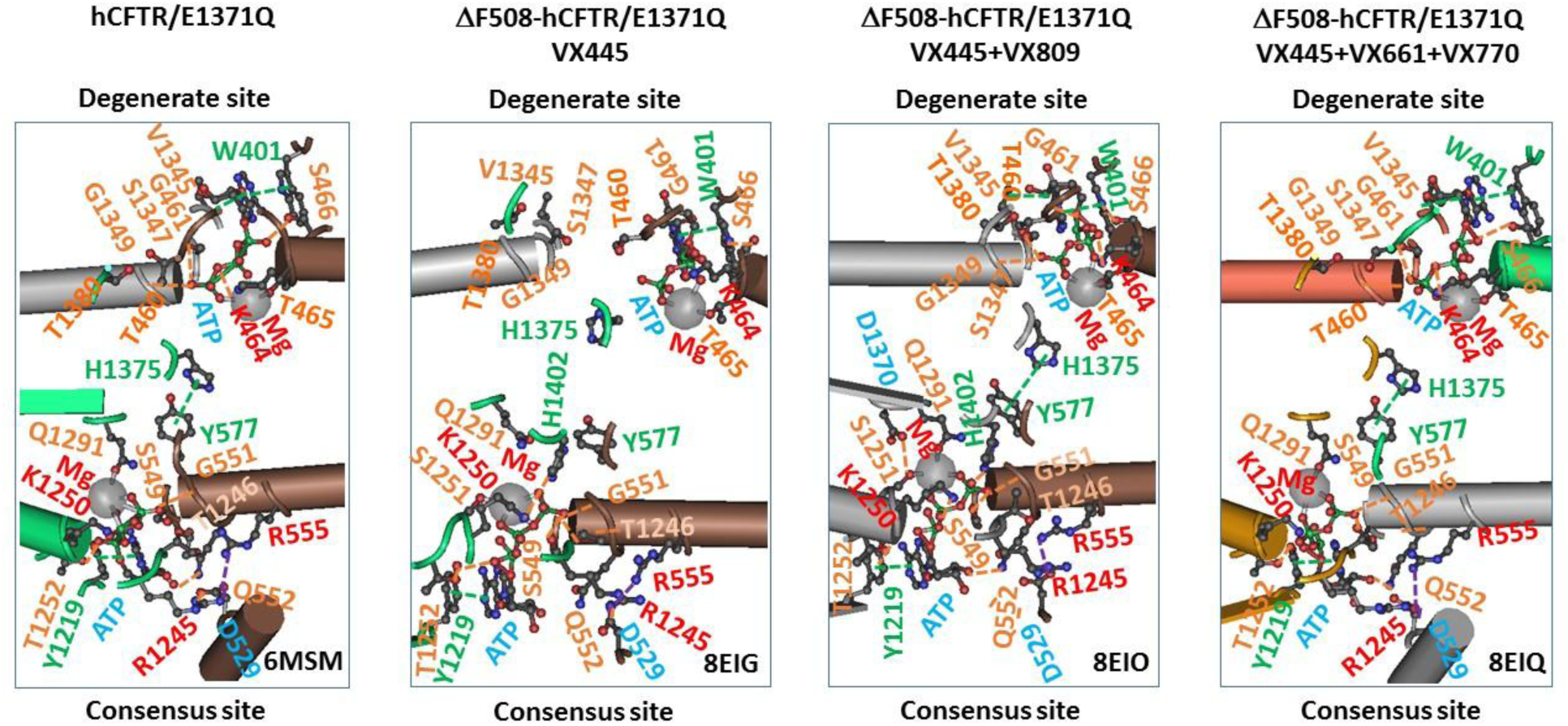
Mg/ATP-mediated NBD1-NBD2 interactions of hCFTR/E1371Q with or without F508-in response to modulators. The cryo-EM structures of CFTR constructs such as open phosphorylated hCFTR/E1371Q with Mg/ATP bound (PDB, 6MSM), open phosphorylated ΔF508-hCFTR-E1371Q with Mg/ATP/Trikafta bound (PDB, 8EIQ), open phosphorylated ΔF508-hCFTR-E1371Q with Mg/ATP/VX-445/VX-809 bound (PDB, 8EIO), closed phosphorylated ΔF508-hCFTR-E1371Q with Mg/ATP/VX-445 bound (PDB, 8EIG) are used for the models.

For example, the D529-R1245 salt bridge or H-bond near the Walker A half-site. More importantly, Y577 from NBD1 formed a strong π-π interaction with H1375 from NBD2. The same Y577-H1375 π-π interaction was also found in ΔF508-hCFTR/E1371Q with Mg/ATP/VX-445/VX-809 bound or hCFTR/E1371Q with Mg/ATP bound. However, the D529-R1245 salt bridge or H-bond and the Y577-H1375 π interaction, together with the traditional interactions between the phosphates and the signature LSHGH half-site, were disrupted in ΔF508-hCFTR/E1371Q with Mg/ATP/VX-445 bound (Figure 4), and the Y577-Y577’ π interaction is not shown in the isolated hCFTR-Δ(RE,RI) homodimers with or without F508 (28). In that regard, it is reasonable that dimerized NBD1 was stabilized along with an increase in the available melting temperature threshold, from 48 °C to 50 °C for hCFTR or from 40 °C to 47 °C for ΔF508-hCFTR no matter whether E1371Q is introduced or RI is deleted or disordered (Table 1).

### 2.5 Partial NBD1-NBD2 dimerization decreases the melting threshold of NBD1

Since the same Y577-H1375 π-π interaction was also missing in the partially dimerized NBD1 and NBD2 of ΔF508-hCFTR/E1371Q with Mg/ATP/VX-445 bound (Figure 4), it is particularly interesting to investigate if the partial NBD dimerization lowers the melting threshold of NBD1.

In response to the removal of lumacaftor and ivacaftor, the salt bridge between K464 and the phosphate group of ATP, and the ATP-S466 H-bond were disconnected (Figure 5a). As a result, the R487-D567 salt bridge, as well as the F446-L454, M472-F490, Y569-M595 and H620-Y625 π interactions were absent. Meanwhile, the T398-K442, R516-K564, S531-K536, D565-Y569 and K606-E608 H-bonds and the K611-F630 and Y625-F626 π interactions were present. Notably, the K503-Y512 π interaction became an H-bond again, along with the shift of H-bonds from K442-S623 and K447-Y627 to K442-S624 and K447-K615 at the interface between N- and C-termini. Collectively, the systematic thermal instability (T_i_) of NBD1 significantly increased from 1.81 in 8EIQ to 2.35 in 8EIG after the total numbers of noncovalent interactions and grid sizes of NBD1 changed from 36 and 65 in 8EIQ to 31 and 73 in 8EIG, respectively (Figure 5a, Table 1).

**Figure 5.**
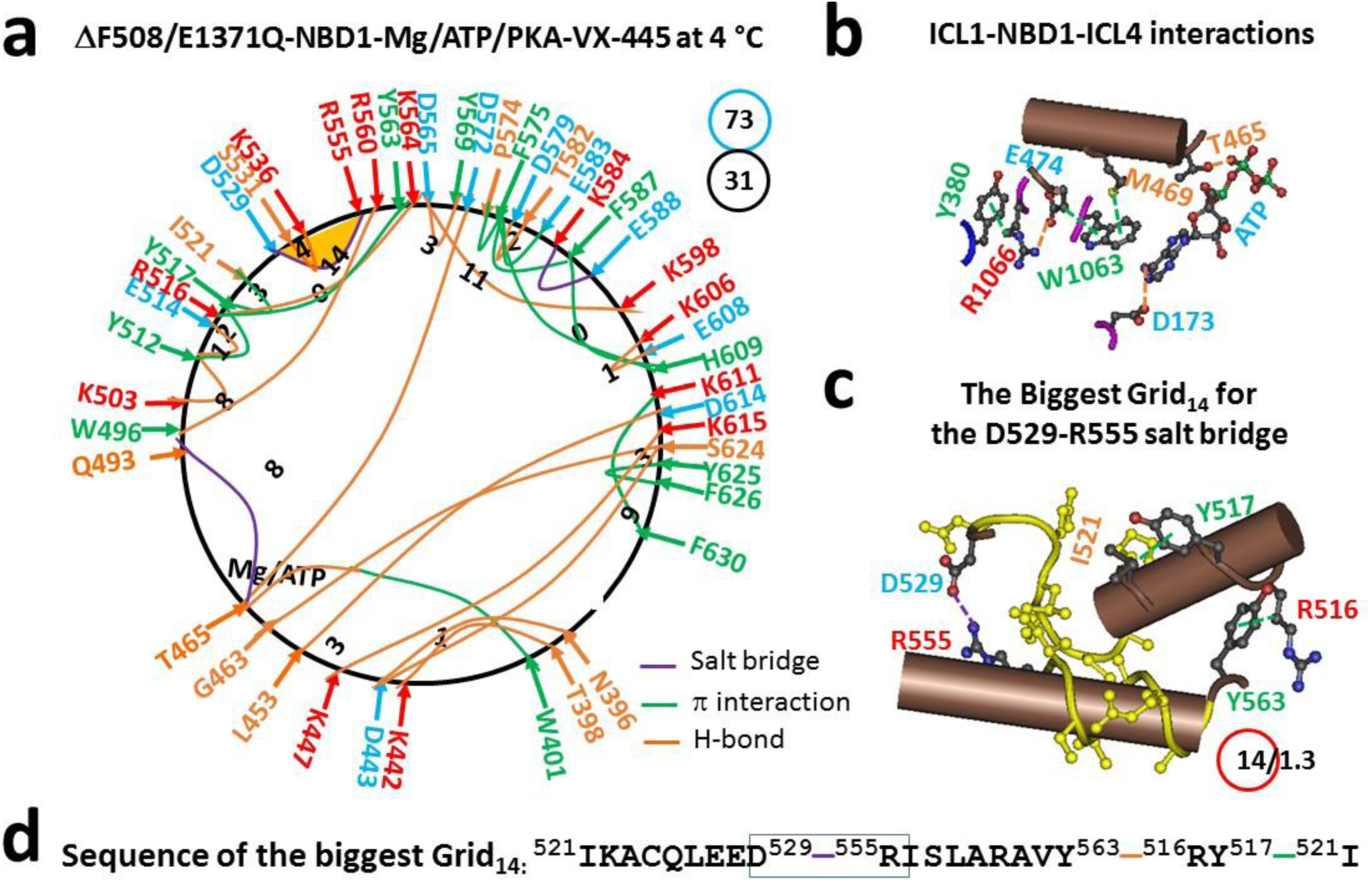
The thermoring structures of NBD1 in phosphorylated hCFTR/E1371Q/ΔF508 with Mg^2+^/ATP/elexacaftor (VX445) bound in the closed state at 4 °C. (**a**) The grid-like noncovalently interacting mesh network of NBD1 based on the cryo-EM structure of hCFTR/E1371Q/ΔF508 with elexacaftor bound in the presence of Mg/ATP/PKA at 4 °C (PDB ID, 8EIG, 3.7 Å). Salt bridges, H-bonds and π interactions are colored purple, orange, and green, respectively. The constrained grid sizes required to control the least-stable noncovalent interactions in the grids are labeled with black numbers. The least-stable D529-R555 salt bridge in the biggest Grid_14_ is highlighted. The total grid sizes and the total grid size-controlled noncovalent interactions along the single peptide chain of NBD1 from G437 to L636 are shown in cyan and black circles, respectively. (**b**) Noncovalent interactions at the ICL1−NBD1-ICL4 interfaces. (**c**) The structure of the biggest Grid_14_ with a 14-residue size to control the least-stable D529-R555 salt bridge. The grid size and the equivalent basic H-bonds for the least-stable noncovalent interaction are shown in and near a red circle. (**d**) The sequence of the biggest Grid_14_ to control the least-stable D529-R555 salt bridge in the blue box.

It is worth noting that the E474-R1066 salt bridge was reestablished as an H-bond at the ICL4-NBD1 interface when ATP connected T465 in NBD1 and D173 in ICL1 (Figure 5b). As a result, the biggest thermoring shifted from Grid_11_ to Grid_14_ in the same α-subdomain to control the weakest D529-R555 salt bridge (Figure 5a). This new thermoring followed a circular path from D529 to I521, Y517, R516, Y563, R555 and back to D529 (Figure 5c-d). With 1.3 basic H-bonds energetically equivalent to this bridge, the calculated T_m,th_ of NBD1 decreased from 47 °C in 8EIQ to 39 °C in 8EIG as expected along with partial NBD1-NBD2 dimerization (Table 1).

### 2.6 Specific fit of the dynamic phosphorylated S813 site to the TMD1/TMD2/NBD1 interfaces primes tight NBD1-NBD2 dimerization

A recent cryo-EM structure revealed that when the PKA catalytic subunit (PKA-C) stabilizes the open state of phosphorylated hCFTR, the phosphate group of S813 is positioned between R487/K564 of NBD1 and R1070 of TMD2 at the NBD1/ICL4 interface (40). However, when the phosphorylated S813 site was introduced into the structure 8EIQ, the phosphate group of S813 was unable to form a salt bridge with R487 and K564 of NBD1 or R1070 of TMD2 in the phosphorylated ΔF508 hCFTR/E1371Q with Mg/ATP/Trikafta bound, which is necessary for tight NBD1-NBD2 dimerization. Instead, it was located between K162 and K166 at ICL1 while Y808 was involved in potential strong π interactions with R1066 at ICL4 and Y380 at the TMD1-NBD1 linker or the distal N-terminal tail of NBD1. Furthermore, R811 H-bonded with E54 and D58 at the N-terminal tail of TMD1. Thus, the phosphorylated S813 site bridged the ICL1/ICL4 interface with the N-terminal tails of TMD1 and NBD1. This bridge was also observed in hCFTR/E1371Q with Mg/ATP bound and ΔF508-hCFTR/E1371Q with Mg/ATP/VX-445/VX-809 bound but not in ΔF508-hCFTR/E1371Q with Mg/ATP/VX-445 bound (Figure 6). This induced fit of the dynamic phosphorylated S813 site to the TMD1/TMD2/NBD1 interfaces of hCFTR/E1371Q may explain why the E54A or D58A or E54A/D58A mutation inhibits the binding of the R domain to the N-terminal tail and the related hCFTR activity (41), and why mutations such as K166A/H, R1066H, S813D/Y808A and S813D/R811A suppress curcumin potentiation of hCFTR activity when curcumin is bound to this S813 binding site (38).

**Figure 6.**
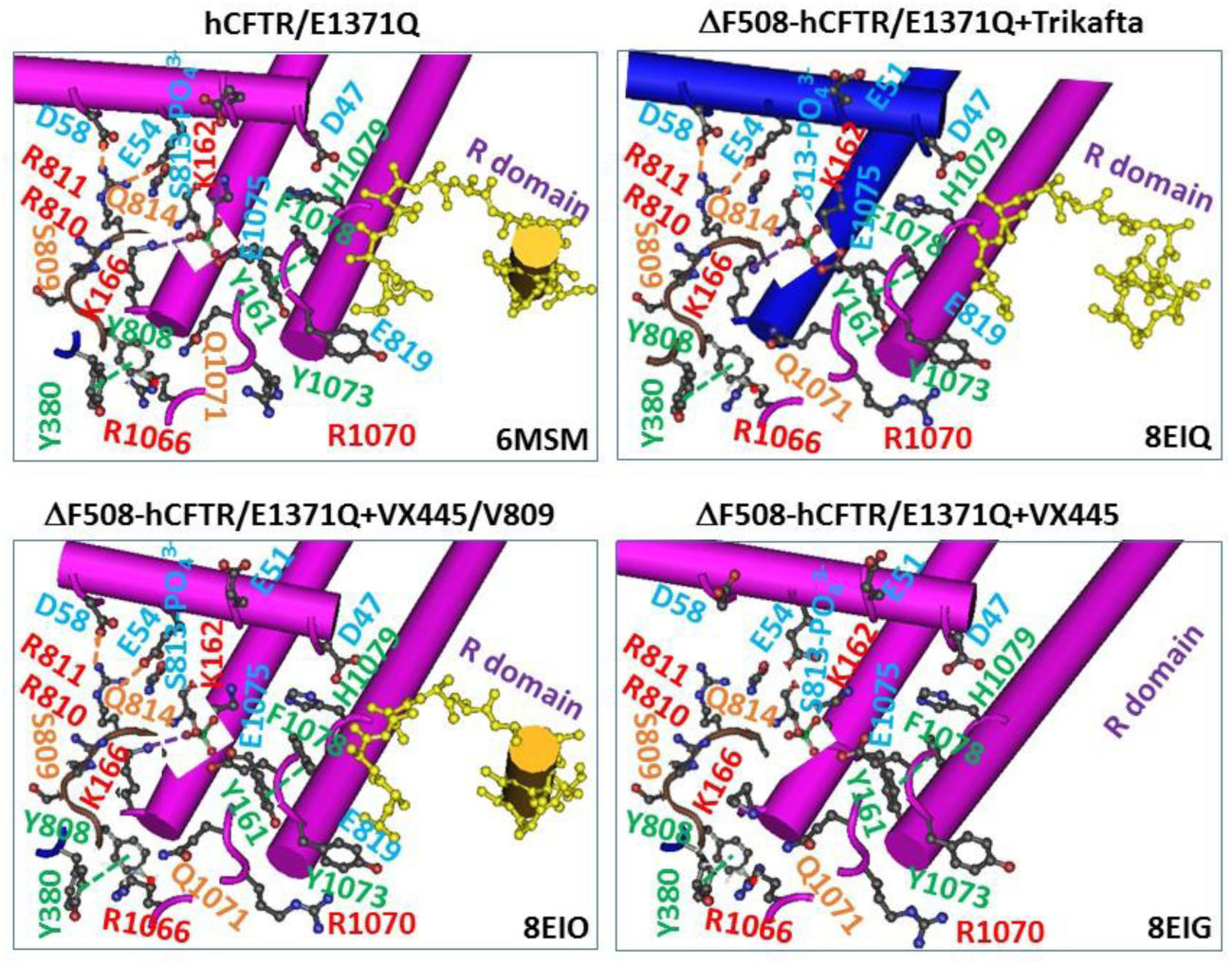
Specific interactions of the dynamic phosphorylated S813 site with ICL1/ICL4/NBD1 of -hCFTR/E1371Q with and without F508 in respons to modulators. The cryo-EM structures of CFTR constructs such as open phosphorylated hCFTR/E1371Q with Mg/ATP bound (PDB, 6MSM), open phosphorylated ΔF508-hCFTR-E1371Q with Mg/ATP/Trikafta bound (PDB, 8EIQ), open phosphorylated ΔF508-hCFTR-E1371Q with Mg/ATP/VX-445/VX-809 bound (PDB, 8EIO), closed phosphorylated ΔF508-hCFTR-E1371Q with Mg/ATP/VX-445 bound (PDB, 8EIG), and open phosphorylated hCFTR-E1371Q with Mg/ATP/PKA-C bound (PDB, 9DW9) are used for the models.

### 2.7 Strong cytoplasmic TMD1-TMD2 interactions promote the specific fit of the dynamic phosphorylated S813 site to the TMD1/TMD2/NBD1 interfaces favoring maximal channel activity

Our previous study showed that K190 and K978 are located in the tetrahelix bundle of hCFTR. Constitutive gain of function (GOF) mutations such as K978C and K190C significantly stimulate ATP-independent but PKA-dependent channel activation, thereby increasing ATP and PKA sensitivity allosterically. Importantly, these constitutive ICL mutations can restore the function of CFTR mutants like G551D without ATP binding and NBD2-free Δ1198 without NBD dimerization (42). They also prevent the irreversible thermal inactivation of ΔF508 hCFTR (10). All of this suggests that the tetrahelix bundle acts as a gating linchpin for ATP-independent but phosphorylation-dependent CFTR activation.

Since the extracellular gate at the selectivity filter remains closed when Y922 (Y914 in hCFTR) is unable to hold the Cl^-^ ion with G104 and T339 (G103 and T338 in hCFTR, respectively) in a cryo-EM structure of zebrafish CFTR (zCFTR) even after R domain phosphorylation and ATP binding allowing tight NBD1-NBD2 dimerization (Figures S1) (39), this structure can serve as an ideal negative control to investigate how the tetrahelix bundle gates the CFTR channel in an ATP-independent but phosphorylation-dependent manner.

In the dephosphorylated closed state of wild type (WT) hCFTR, the segment 970-985 in TM9 was structured as an α-helix and connected with TM4 via the R251-D985 H-bond. R251 and S256 in TM4 were located on two separate α-helices. Similarly, T1053 and E1046 in TM10 were also positioned on two separate α-helices. The C-terminal region of the R domain was wedged among TMs 3, 4, 9, and 10 (Figure 7). This is reminiscent of the inhibitory lipid at the active vanilloid site in heat-gated transient receptor potential vanilloid-1 (TRPV1) or 3 (TRPV3) bio-thermometers (33–34).

**Figure 7.**
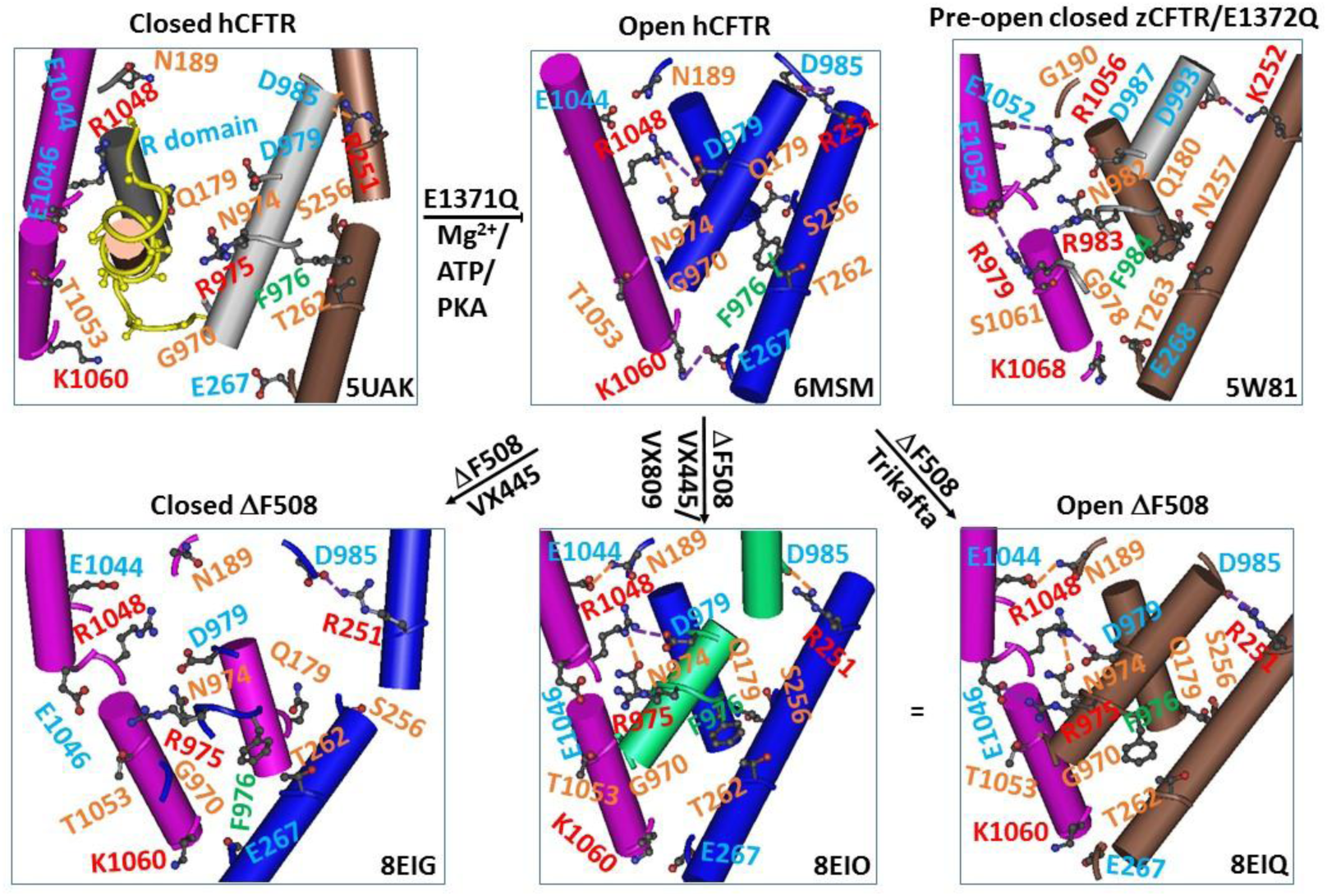
Phosphorylation-dependent cytoplasmic TMD1-TMD2 interactions of hCFTR/E1371Q with or without F508 in repsonse to modulators. Closed and open hCFTR and pre-open closed zCFTR are controls. The cryo-EM structures of CFTR constructs such as closed dephosphorylated hCFTR (PDB, 5UAK), open phosphorylated hCFTR-E1371Q with Mg/ATP bound (PDB, 6MSM), pre-open closed phosphorylated zCFTR-E1372Q with Mg/ATP bound (PDB, 5W81), open phosphorylated ΔF508-hCFTR-E1371Q with Mg/ATP/Trikafta bound (PDB, 8EIQ), open phosphorylated ΔF508-hCFTR-E1371Q with Mg/ATP/VX-445/VX-809 bound (PDB, 8EIO) and closed phosphorylated ΔF508-hCFTR-E1371Q with Mg/ATP/VX-445 bound (PDB, 8EIG) are used for the models.

When phosphorylation released the R domain for channel opening, both TM4 and TM10 underwent a π-to-α transition. In addition to the R251-D985 salt bridge, R1048 also formed an H-bond or a salt bridge with N974 or D979 via their side chains, leading to a smaller Grid_4_. Meanwhile, F976 formed a CH-π interaction with T262 via their side chains. Notably, K1060 linked with E267 via a salt bridge, confirming the previous report (Figure 7) (43).

In contrast, when exposed to Mg/ATP agonists that stimulate the phosphorylated zCFTR, the loose α-helix in TM9 extended to D987 (D979 in hCFTR), and segments 1052-1056 and 1057-K1068 in TM10 formed two α-helices. In this case, most noncovalent interactions were disrupted, including the D987-R1056-N982 (D979-R1048-N974 in hCFTR) salt bridge or H-bond and the F984-T263 (F976-T262 in hCFTR) π interaction. Instead, only the E1052-R1056 and R979-E1054 salt bridges were present in this pre-open closed state (Figures 4 and S2). Therefore, the tight cytoplasmic TMD1-TMD2 interactions were required for CFTR opening.

Since there is no gap between TMD1 and TMD2 of phosphorylated ΔF508-hCFTR upon Trikafta and Mg/ATP binding (22), it is necessary to examine if Trikafta tightens the cytoplasmic TMD1-TMD2 interactions. The study on the tertiary structure of this activated mutant indicated that despite the disruption of the F976-T262 π interaction and the E267-K1060 salt bridge, TM10 exhibited two α−helices at the T1053-E1046 interface. This allowed for a new E1044-N189 H-bond and a new R975-E1046 salt bridge at the TM9-TM10-TM3 interfaces. Thus, a much smaller Grid_1_ produced a more thermally stable triangle among N974, R975, E1046, and R1048 (Figure 7). Since VX-809 alone stimulates ΔF508-CFTR activity in the absence of tight NBD dimerization (22, 26), the gating defect of ΔF508-hCFTR was first rescued upon the tight cytoplasmic TMD1-TMD2 interactions before the thermal defect of NBD1 was corrected.

Further detailed studies showed that all three modulators in Trikafta worked together to enhance this unique cytoplasmic H-bond network through their specific noncovalent interactions with TMD1 and TMD2 (Figure S3), which were similar to previous reports (22, 44–46). The same H-bond/salt bridge network was also observed in ΔF508-hCFTR/E1371Q with Mg/ATP/VX-445/VX-809 bound for tight NBD dimerization. However, it was compromised in ΔF508-hCFTR/E1371Q with Mg/ATP/VX-445 bound for partial NBD1-NBD2 dimerization (Figures 4 & 7). Thus, once Trikafta binding tightened the cytoplasmic TMD1-TMD2 interactions, the dynamic phosphorylated S813 site could fit into the ICL1/ICL4/NBD1 interfaces for the tight NBD1-NBD2 interactions.

## 3. DISCUSSION

F508 in NBD1, like F251 in β-D-glucoside glucohydrolase (46), plays a central role in CFTR expression on the cell surface and ATP/PKA-mediated channel activity. The deletion of F508 (ΔF508) on the surface of NBD1 destabilizes NBD1 and the nearby interdomain interactions, leading to both thermal and gating defects (17–21). Thus, rescuing strategies have focused on this domain and its interfaces (20, 47–51). Although Trikafta modulators do not target them, they successfully correct both defects of phosphorylated ΔF508-CFTR even at 37 °C along a significantly different pathway (23–27). This study demonstrated that Trikafta binding to flexible TMD1 and TMD2 stabilized the most flexible NBD1 by reinforcing specific phosphorylation-dependent interdomain interactions, rescuing defects of F508del with improved channel opening and chloride ion transport. These gating features are vital for maintaining airway surface liquid homeostasis and overall lung function. Although the deletion of F508 causes the primary thermal defect in NBD1 and then the gating defect at the TMD1/TMD2 interface, Trikafta rescued them in a reverse manner allosterically.

### 3.1 Specific binding of the dynamic phosphorylated S813 site to the TMD1/TMD2/NBD1 interfaces is necessary for the normal gating pathway

Mg/ATP-mediated NBD1-NBD2 dimerization is required for the normal channel opening of phosphorylated CFTR (14–16). However, recent simultaneous kinetic studies of NBD isomerization and channel gating have shown that CFTR quickly transitions to a flicker-closed state upon NBD dimerization (52). Consistent with this finding, zCFTR remains closed even when NBD1 and NBD2 are tightly dimerized by Mg/ATP binding at their interface (39).

On the other hand, our previous studies have highlighted the significance of constitutive mutations K190C and K978C at the TMD1/TMD2 interface in ATP-independent but phosphorylation-dependent hCFTR activation (42), as well as the significance of the E267-K1060 salt-bridge at this interface in allosterically coupling to R domain phosphorylation (43). This study further suggests that tight cytoplasmic TMD1-TMD2 interactions are crucial for CFTR opening and are linked to specific interactions of the dynamic phosphorylated S813 site with ICL1, ICL4 and the distal N-terminal tails of TMD1 and NBD1 (Figures 6-8).

**Figure 8.**
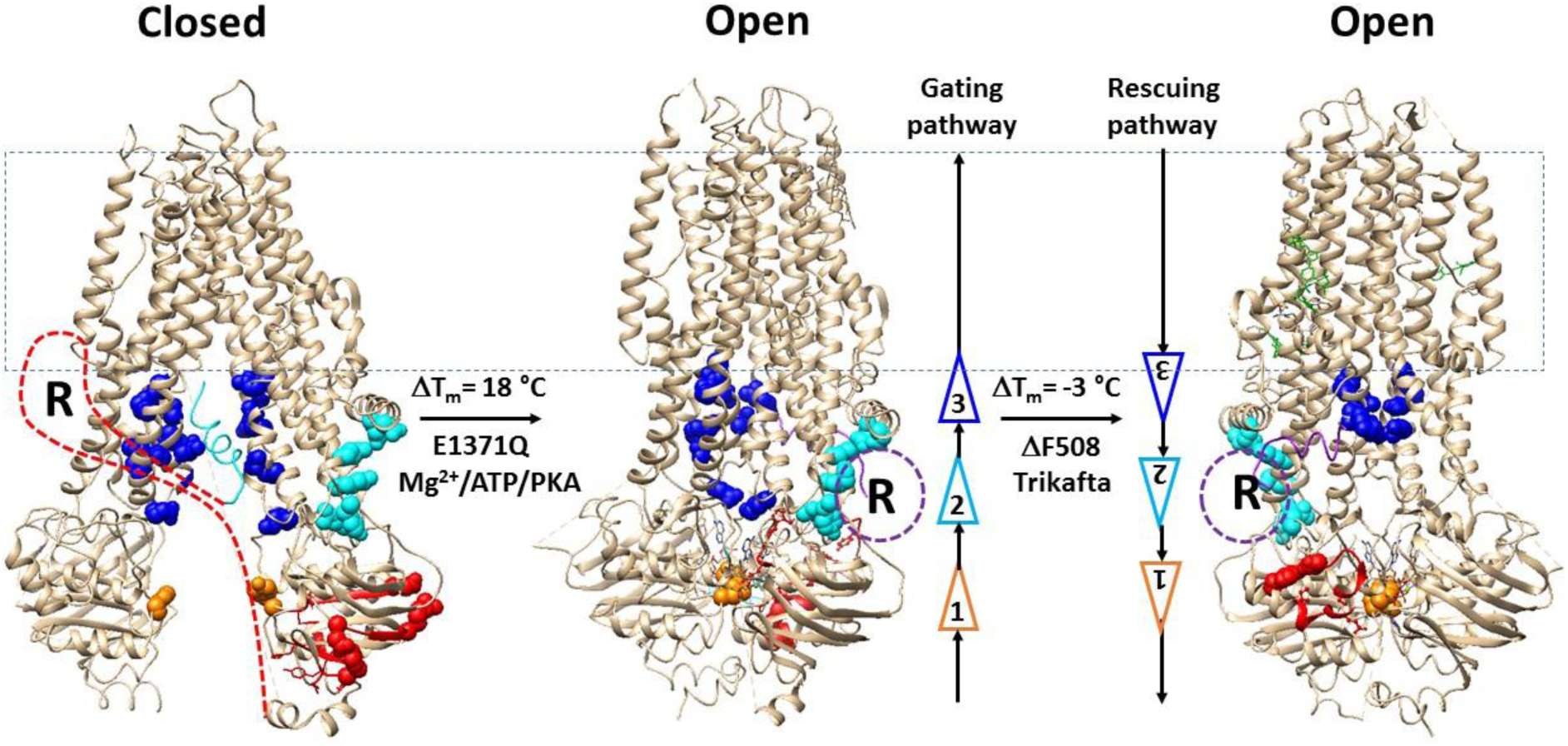
Putative pathway for Trikafta to correct the primary gating and subsequent thermal defects of ΔF508-CFTR. Cryo-EM structures of closed dephosphorylated hCFTR (PDB: 5UAK), open phosphorylated hCFTR/E1371Q with Mg/ATP bound (PDB: 6MSM) and open phosphorylated hCFTR/ΔF508/E1371Q with Mg/ATP/Trikafta bound (PDB: 8EIQ) are used for the models. The red curves and the purple circles represent the dephosphorylated and phosphorylated R domain, respectively. The critical residues for cytoplasmic TMD1-TMD2 interactions are colored blue. The C-terminal region of the R domain and S813-binding sites are colored in cyan. The residues for the NBD dimerization hallmark, the Y577-H1375 π bridge, are colored in orange. Three modulators such as VX-445, VX-661 andVX-770 are colored green and the biggest thermorings to control the least-stable noncovalent interactions in NBD1 are shown in red. The residues responsible for the least-stable noncovalent interactions in NBD1 are shown in space fills. Three key interdomain interactions are shown as triangles.

Moreover, the phosphorylated R domain has been found near the ICL1-ICL4 interface of not only the open E1371Q-hCFTR but also the glutathione transporter Yeast Cadmium Factor 1 (Ycf1) E1435Q mutant (16, 53), Therefore, the flexible cytoplasmic TMD1-TMD2 interface cannot guarantee strict coupling between NBD dimerization and channel opening. Specific binding of the dynamic phosphorylated S813 site to the TMD1/TMD2/NBD1 interfaces is necessary to enhance cytoplasmic TMD1-TMD2 interactions for normal channel opening (Figure 8).

### 3.2 Specific binding of the dynamic phosphorylated S813 site to the TMD1/TMD2/NBD1 interfaces is also necessary for the Trikafta correction pathway

This study further indicated that despite the weakened NBD1-ICL4 interactions in ΔF508 CFTR, the tight cytoplasmic TMD1-TMD2 interactions in ΔF508 CFTR/E1371Q were not restored by VX-445 until the combination of VX-445 and VX-809 or Trikafta (VX-445, VX-661 and VX-770) were bound to TMD1 and TMD2 (Figure 7). Along with these interactions, the specific interactions of the dynamic phosphorylated S813 site with ICL1, ICL4 and the distal N-terminal tails of TMD1 and NBD1 and Mg/ATP-mediated tight NBD1-NBD2 dimerization were also observed in ΔF508 CFTR/E1371Q with VX-445 and VX-809 or Trikafta bound rather than VX-445 alone bound (Figures 4, 6-7). Therefore, similar to lysine ligation to the heme iron in cytochrome c (54), remote Trikafta binding to flexible regions of TMD1 and TMD2 may reinforce the cytoplasmic TMD1-TMD2 interactions, favoring the specific binding of the dynamic phosphorylated S813 site to the TMD1/TMD2/NBD1 interfaces and NBD dimerization that further stop irreversible thermal inactivation by stabilizing NBD1 (Figure 8) (10, 12, 37). Based on this important role of the cytoplasmic TMD1/TMD2 interactions, it is reasonable to find some CF-causing mutations G970E/D, D979V, E1044X, E1046X, and T1053I are involved (http://www.genet.sickkids.on.ca/cftr; https://cftr2.org).

### 3.3 Role of a curcumin-like drug in optimizing cystic fibrosis treatment

Curcumin can strengthen the stimulatory interactions between the dynamic phosphorylated S813 site and the ICL1/ICL4 interface to stabilize the open state (38, 55). However, curcumin can also stabilize the stochastic, competitive and stimulatory binding of the phosphorylated S795 site to these interfaces (38). This may explain why curcumin alone is not effective in maintaining ΔF508 CFTR activity at room temperature (56), or preventing the irreversible thermal inactivation of ΔF508-hCFTR at physiological temperature (10).

On the other hand, Trikafta modulators maintain higher ΔF508-hCFTR activity even at a physiological temperature of 37 °C (23–27). Therefore, specific interactions of the dynamic phosphorylated S813 site with not only the ICL1-ICL4 interface but also the distal N-terminal tails of TMD1 and NBD1 may be necessary to couple tight cytoplasmic TMD1-TMD2 interactions to Mg/ATP-mediated NBD1-NBD2dimerization at human body temperature. In support of this proposal, when a strong water-like spherical density at the NBD1/TMD1/TMD2 interfaces was observed at a low temperature of 4 °C (22), more protein-water H-bonds were also reported in the cold-denatured state of yeast frataxin than in the hot-denatured state (57). Thus, although a low temperature may hydrate the NBD1/TMD1/TMD2 interfaces through an interfacial H-bonding network between the ordered water molecules and the charged residues such as K162, K166, E54, D58 or R1066, a high temperature may dehydrate these interfaces, allowing specific binding of the dynamic phosphorylated S813 site to the NBD1/TMD1/TMD2 interfaces to be captured in a cryo-EM structure at a physiological temperature. Most importantly, once this specific binding can be stabilized by a drug like curcumin, a combined therapy of Trikafta with this drug may further optimize cystic fibrosis treatment through improved protein durability and drug efficacy. Further molecular dynamics (MD) simulations are necessary to accurately understand the role of the dynamic binding of the phosphorylated S813 site to the NBD1/TMD1/TMD2 interfaces in the normal gating pathway of native CFTR, as well as in the correction pathway of ΔF508-CFTR by Trikafta.

## 4. CONCLUSIONS

The ability of combination therapy to correct the thermal and gating defects of misfolded CFTR is essential for the optimal treatment of cystic fibrosis. While Trikafta modulators have been approved to treat the most common cystic fibrosis mutation F508del-CFTR, less is known about the exact rescuing pathway. This study demonstrated that Trikafta binding can correct functional defects by enhancing specific phosphorylation-dependent interdomain interactions to stabilize the most flexible domain in the mutant allosterically. These findings highlight the importance of post-translational chemical modifications in protein function. Targeting this correction pathway could be a promising approach for developing new therapies for patients with CF who have similar or different CFTR mutations.

## 5. COMPUTATIONAL METHODS

### 5.1 Data mining resources

Four cryo-EM structures of human CFTR (hCFTR) constructs with or without the F508 deletion at 4 **°**C were selected for thermoring analysis. These structures included dephosphorylated hCFTR in a closed state (PDB ID, 5UAK, model resolution = 3.87 Å) (14), phosphorylated E1371Q with Mg/ATP bound in an open state (PDB ID, 6MSM, model resolution = 3.2 Å) (16), phosphorylated hCFTR/E1371Q/ΔF508 with Mg/ATP/elexacaftor bound in a closed state (PDB ID, 8EIG, model resolution = 3.7 Å), and phosphorylated hCFTR/E1371Q/ΔF508 with Mg/ATP/Trikafta bound in an open state (PDB ID, 8EIQ, model resolution = 3.0 Å) (22). Additionaly, the cryo-EM structure of zebrafish CFTR (zCFTR) construct with phosphorylation and Mg/ATP bound (PDB, 5W81, model resolutions 3.4 Å) was used as a control to demonstrate the role of the cyroplasmic TMD1/TMD2 interface (39). Finally, the cryo-EM structure of phosphorylated hCFTR with Mg/ATP/PKA-C bound (PDB, 9DW9, model resolution = 2.8 Å) was also used to fit the dynamic phosphorylated S813 site to the NBD1/TMD1/TMD2 interfaces (40).

### 5.2 Standard methods for filtering non-covalent interactions

The standard methods for filtering non-covalent interactions, along with exact calculations, were the same as those previously used, ensuring accurate and repeatable results (30–37). UCSF Chimera was used to review potential stereo-selective or regio-selective intra-domain lateral noncovalent interactions along the single peptide chain of hNBD1 with or without F508. These interactions included salt bridges, H-bonds and lone pair/CH/cation-π interactions between paired amino acid side chains. When a backbone NH or CO at a residue position was within a cutoff distance for an H-bond with a side chain of a nearby residue, that H-bond was also considered. Detailed cutoff distances and interaction angles can be obtained in the online Supporting Information (Tables S1, S2, and S3). In this study, approximately at least 18 different noncovalent interactions were identified along the single peptide chain from L383 to L644 of NBD1 on each protomer.

### 5.3 Mapping the thermoring structures of NBD1 using the grid thermodynamic model

The same grid thermodynamic model that was previously validated was used to map the thermoring structures of NBD1 in full-length CFTR (30–37). Briefly, filtered noncovalent interactions were placed in an adjacency matrix to create a systematic fluidic grid-like mesh network along the single peptide chain (black line) of NBD1 from L383 to L644. In this network, each node represented paired protein residues (colorful arrows) that formed a specific noncovalent interaction via their side chains, and each colorful edge represented a specific noncovalent interaction that formed a direct path. However, a reverse path represented a sequence of alternating peptide chain and links with no repeating nodes. In these representations, any link between two nodes had two path lengths that indicated the cost of traveling across the link. In this study, the path length was defined as the total number of free or silent amino acids that did not involve any noncovalent interactions between two nodes. Thus, the traveling cost across any link was determined by the reverse path length because the direct path length was zero, and any noncovalent inteaction with the alternating peptide chain lining a reverse path had a positively curved edge (58). For example, when F508 formed a π interaction with R560 in Figure 1a, if a direct path started from F508 and ended at R560, the peptide segment from R560 to F508 paved a reverse path.

When multiple reverse paths exist for a specific noncovalent interaction, identifying the shortest reverse path is essential to minimize travel costs. This is achieved by Floyd-Warshall’s Algorithm, which compares all possible adjacent reverse paths through the graph between each pair of nodes (59). In the case of the F508-R560 π interaction shown in Figure 1a, one adjacent reverse path from R560 to R553, S495, and F508 had a length of 18 (the total number of free amino acids in peptide segments 507 to 494 and 552 to 559). However, another adjacent reverse path from R560 to Y563, Y517, Y512, and F508 included Y512-Y517-Y563 π interactions, resulting in a length of 5 (the sum of free amino acids in peptide segments 509 to 511 and 562 to 561). Since no other possible adjacent reverse paths were shorter than 5, the shortest reverse path length for the F508-R560 bridge was determined to be 5.

Once the shortest reverse path was identified for each noncovalent interaction, the direct path and the shortest reverse path emerged as a thermosensitive ring or “thermoringˮ to control the heat unfolding of that noncovalent interaction. Since the shortest reverse path length was defined as the size of a thermoring, the thermoring was denoted as Grid_s_. In this way, all noncovalent interactions could be tracked by their respective unshared thermoring sizes so that the biggest thermoring could be identified. As the least-stable noncovalent interaction in the biggest thermoring was typically the weakest link along the single peptide chain, a melting temperature threshold (T_m,th_) of NBD1 could be calculated theoretically. Furthermore, when the total non-covalent interactions and grid sizes along the given polypeptide chain were displayed by black and cyan circles beside the mesh network map, the grid-based systematic thermal instability (T_i_) could also be computed to evaluate the flexibility of NBD1.

### 5.4 Calculation of the melting temperature threshold (T_m,th_)

The same empirical equation and coefficients used in previous studies on temperature-dependent structures were applied to calculate the melting temperature thresholds (T_m,th_) for thermal unfolding of a specific grid (30–37):

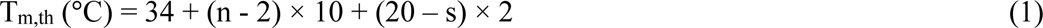

where, n represents the total number of basic H-bonds (each approximately 1 kcal/mol) that are calculated to be approximately equal in stability of the least-stable noncovalent interaction controlled by the given grid (60); and s is the grid size used to control the least-stable noncovalent interaction in the given grid.

### 5.5 Evaluation of the grid-based systemic thermal instability (T_i_)

The same empirical equation used in previous studies on temperature-dependent structures was utilized to calculate the systematic thermal instability (T_i_) along the given polypeptide chain (30–37):

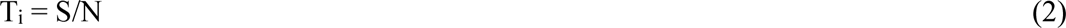

where, S and N are the total grid sizes and the total non-covalent interactions along a specific polypeptide chain. This calculation allows for evaluation of the protein’s compact conformational entropy or flexibility.

## AUTHOR CONTRIBUTIONS

Guangyu Wang wrote the main manuscript text and prepared Figures. 1, 2, 3, 4, 5, 6, 7, 8 and Table 1. Supporting Information (Tables S1, S2, S3, S4; Figures S1, S2, S3) and reviewed the manuscript.

## ACKNOWLEDGEMENTS

The author’s own studies cited in this article were supported by the NIDDK Grant (DK45880 to D.C.D.) and the Cystic Fibrosis Foundation grant (DAWSON0210), the NIDDK grant (2R56DK056796-10) and the American Heart Association (AHA) Grant (10SDG4120011 to GW).

## ETHICS STATEMENT

The author confirms that he has followed the ethical policies of the journal.

## DATA AVAILABILITY STATEMENT

All data generated or analyzed during this study are included in this published article and Supporting Information.

## SUPPLEMENTARY INFORMATION

Additional supporting information can be found online in the Supporting Information section at the end of this article.

## ABBREVIATIONS

ABC: ATP-binding cassette
CF: cystic fibrosis
cryo-EM: cryoelectron microscopy
CFTR: cystic fibrosis transmembrane conductance regulator
DSC: differential scanning calorimetry
DSF: differential scanning fluorimetry
GOF: gain of function
hCFTR: human
CFTR ICL1: intracellular loop 1
ICL4: intracellular loop 4
MD: molecular dynamics
NBD1: nucleotide binding domain 1
NBD2: nucleotide binding domain 2
PKA: protein kinase A
PKA-C: the PKA catalytic subunit
R: regulatory
RE: regulatory extension
RI: regulatory insert
T_i_: systematic thermal instability
T_m,th_: melting temperature threshold
TMH: transmembrane helix
TMD1: transmembrane domain 1
TMD2: transmembrane domain 2
TRPV1: transient receptor potential vanilloid-1
TRPV3: transient receptor potential vanilloid-3
2-APB: 2-aminoethoxydiphenyl borate
WT: wild type
wtCFTR: wild type CFTR
Ycf1: Yeast Cadmium Factor 1
zCFTR: zebrafish CFTR

