## Supplemental Tables 1-4 and Figures S1-S3 for "Trikafta rescues F508del-CFTR by tightening specific phosphorylation-dependent interdomain interactions"

Short title: Correction pathway of ΔF508-CFTR

Guangyu Wang 1, 2*

[1]Department of Physiology and Membrane Biology, University of California School of Medicine, Davis, California, USA

[2]Department of Drug Research and Development, Institute of Biophysical Medico-chemistry, Reno, Nevada, USA

**This Supporting Informational material includes:**
**Tables S1, S2, S3 and S4, and Figures S1, S2 and S3**

**Table S1 Noncovalent interactions along the single peptide chain of NBD1 from E384 to M645 without MgATP-bound in hCFTR subunit at 4 °C (PDB ID, 5UAK)**

| **Noncovalent interaction** | **Cut-off distance** | **Linked residues** |
|---|---|---|
| Salt/metal bridge | 3.2-4 Å | R450-D567, D529-K532, K584-E608-K612 |
| H-bond | <3.9 Å donor-H-acceptor <60° | D443-Y627, R487-D567/L568, S495-R553, **D579-T582**, E583-K606, T629-E632 |
| π–π interaction | 2.65–6.5 Å | **Y512-Y517**-Y563, **F575-F587-H609,** H620-Y625 |
| cation–π interaction | <6.0 Å | |
| $CH_3$/CH–π interaction | 2.65-3.01 Å | F508-R560, F640-L644 |
| Lone pair–π interaction | 3-3.7 Å | |

Note: Bold interactions were shared between hCFTR and hCFTR/E1371Q+Mg/ATP.

**Table S2. Noncovalent interactions along the single peptide chain of NBD1 from E384 to Q637 with MgATP-bound in hCFTR/E1371Q subunit at 4 °C (PDB ID, 6MSM)**

| **Noncovalent interaction** | **Cut-off distance** | **Linked residues** |
|---|---|---|
| Salt/metal bridge | 3.2-4 Å | **T465/Q493-$Mg^{2+}$,** K503-D537, **D529-K555** |
| H-bond | <3.9 Å donor-H-acceptor <60° | E391-K447-Y627, T398-K442, G463-E621, K464/**T465**/S466/Q493-**ATP**, **T465-D572**, W496-R560, **Y512-E514**, **Y517-D537**, Q525-E585, **D529-Q552**, **D579-T582**, E583-K606, T629-S631 |
| π–π interaction | 2.65–6.5 Å | F400-F409, **W401-ATP**, W496-F508, **Y512-Y517**, **F575-H609-F587-F575** |
| cation–π interaction | <6.0 Å | K503-Y512 |
| $CH_3$/CH–π interaction | 2.65-3.01 Å | M394-F446-L454, **W401-L475**, M472-F490, F508-R560, **Y517-I521**, F533-I539, **P574-F575,** F626-L633 |
| Lone pair–π interaction | 3-3.7 Å | Y569-M595, F587-C592 |

Note: Bold interactions were shared between hCFTR/E1371Q and hCFTR/E1371Q/ ΔF508 with Mg/ATP bound.

**Table S3. Noncovalent interactions along the single peptide chain of NBD1 from L383 to L636 with MgATP-bound in hCFTR/E1371Q/ΔF508 subunit with Trikafta bound at 4 °C (PDB ID, 8EIQ)**

| Noncovalent interaction | Cut-off distance | Linked residues |
|---|---|---|
| Salt/metal bridge | 3.2-4 Å | **T465-$Mg^{2+}$-Q493**, R487-D567, **D529-K555**, D565-K598, E608-K611 |
| H-bond | <3.9 Å donor-H-acceptor <60° | N396-D443, W401-S466, K442-S623, K447-Y627, L453-D614, **G463-E621**, K464/**T465**/S466-**ATP**, **T465-D572**, W496-R560, **Y512-E514**, **Y517-D537**, E528-S531, **D529-Q552**, **D579-T582**, K584-E588 |
| π–π interaction | 2.65–6.5 Å | **W401-ATP**, **Y512-Y517**, **F575-H609-F587-F575,** H620-Y625 |
| cation–π interaction | <6.0 Å | K503-Y512, R516-Y563 |
| $CH_3$/CH–π interaction | 2.65-3.01 Å | **W401-L475**, F446-L454, M472-F490, **Y517-I521**, **P574-F575** |
| Lone pair–π interaction | 3-3.7 Å | Y569-M595 |

Note: Bold interactions were shared between hCFTR/E1371Q and hCFTR/E1371Q/ ΔF508 with Mg/ATP bound.

**Table S4. Noncovalent interactions along the single peptide chain of NBD1 from G437 to L636 in phosphorylated hCFTR/E1371Q/ΔF508 with $Mg^{2+}$/ATP/elexacaftor (VX445) bound in the closed state at 4 °C (PDB ID, 8EIG).**

| **Noncovalent interaction** | **Cut-off distance** | **Linked residues** |
|---|---|---|
| Salt/metal bridge | 3.2-4 Å | **T465/Q493-$Mg^{2+}$ , D529-K555**, K584-588 |
| H-bond | <3.9 Å donor-H-acceptor <60° | N396-D443-S624-G463, T398-K442, K447-K615, L453-D614, **ATP-T465-D572**, **W496-R560**, K503-**Y512-E514**, R516-K564, S531-K536, D565-Y569/K598, **D579-T582,** K606-E608 |
| π–π interaction | 2.65–6.5 Å | **W401-ATP**, **Y512-Y517**, **F575-H609-F587-F575,** Y625-F626 |
| cation–π interaction | <6.0 Å | R516-Y563 |
| $CH_3$/CH–π interaction | 2.65-3.01 Å | **Y517-I521**, **P574-F575**, F575-E583, K611-F630 |
| Lone pair–π interaction | 3-3.7 Å | |

Note: Bold interactions were shared between hCFTR/E1371Q and hCFTR/E1371Q/ΔF508 with Mg/ATP bound.

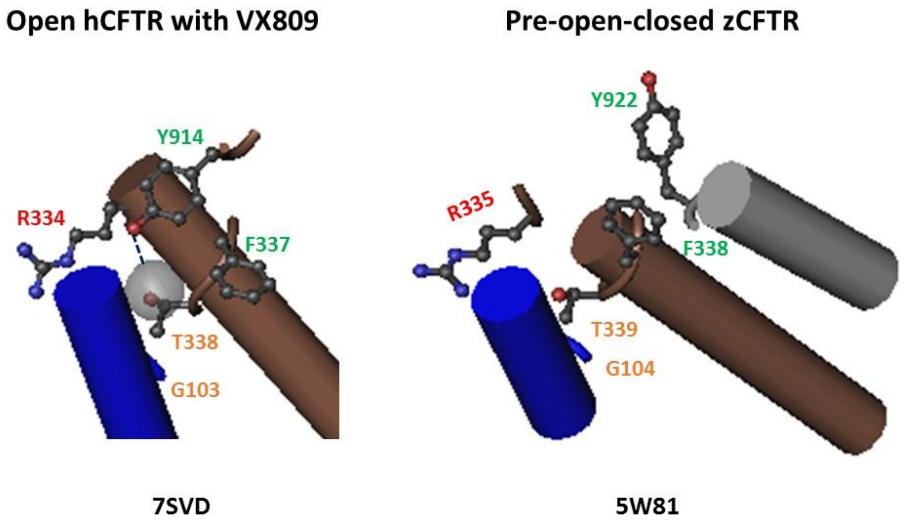


**Figure S1**. Comparison between pre-open closed zCFTR and open hCFTR with VX-809 bound at the extracellular gate.

| | TM3 | | TM4 | | | | TM9 | TM10 |
|---|---|---|---|---|---|---|---|---|
| | 179 | 189 | 251 | 256 | 262 | 267 | 970 974 979 985 | 1044 1048 1053 1060 |
| hCFTR | Q | N | R | S | T | E | GGILNRFSKDIAILDD | ESEGRSPIFTHLVTSLK |
| zCFTR | Q | G | K | T | N | E | GRIMNRFTKDMATIDD | ETEARSPIFSHLIMSLK |
| | 180 | 190 | 252 | 257 | 263 | 268 | 978 982 987 993 | 1052 1056 1061 1068 |

**Figure S2**. Sequence alignment of hCFTR and zCFTR

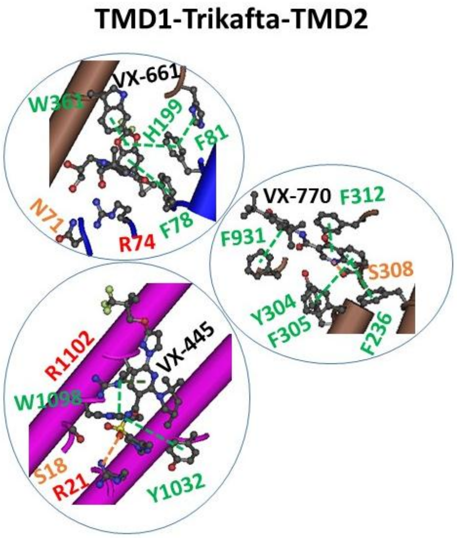


**Figure S3**. The binding sites of three modulators (VX-445, VX-661 and VX-770) around the TMD1-TMD interface.
